# Insulin docking within the open hemichannel of connexin 43 may reduce risk of amyotrophic lateral sclerosis

**DOI:** 10.1101/2022.05.06.490928

**Authors:** Steven Lehrer, Peter H. Rheinstein

## Abstract

**Background:** Type 2 diabetes (T2D), characterized by hyperinsulinemia, protects motor neurons against amyotrophic lateral sclerosis (ALS). Type 1 diabetes and a total lack of insulin are associated with increased risk of ALS. Connexin 43 (Cx43), an astrocyte protein, operates as an open pore via which toxic substances from the astrocytes reach the motor neurons.

**Methods:** In the current study we performed molecular docking of insulin with monomeric Cx31, monomeric Cx43, and hexameric Cx31 to assess whether insulin might affect the pore. Hexameric Cx31 and hexameric Cx43 have hemichannels composed of 6 subunits that work as transmembrane channels, binding together to form gap junction intercellular channels that open and close. We used the program AutoDock Vina Extended for the molecular docking study.

**Results:** Cx31 shares amino acid and structural similarity to Cx43 and insulin docks to the same position of the N-terminal domain of monomeric Cx31 and monomeric Cx43. Insulin docks within the open hemichannel of hexameric Cx31, potentially blocking it. The block may be responsible for the protective relationship of T2D to ALS.

**Conclusion:** Insulin, especially intranasal insulin, might be a treatment for ALS. An insulin secretogogue such as oral sulfonylurea or glinide might also be of value.

---

Amyotrophic lateral sclerosis (ALS) is a disease of motor neurons that affects up to 30,000 people in the United States each year, with 5,000 new cases being diagnosed. Muscles become weaker over time, affecting physical function, and eventually leading to death. The condition has no single cause and no recognized remedy.

Type 2 diabetes (T2D), but not type 1, protected against ALS in a Danish population-based study [1]. A Swedish population study identified a significant inverse association between ALS and T2D, but not type 1 diabetes, with the strongest inverse association 6 years after diabetes onset [2]. An Italian cohort study revealed a significantly reduced ALS risk in T2D (hazard ratio 0.30) with no effect of gender, age, or ALS phenotype [3]. Zhang et al reported that genetically predicted T2D was associated with significantly lower odds of ALS both in European and East Asian populations [4]. Type 1 diabetes, characterized by a total lack of insulin, is associated with increased risk of ALS [1, 2].

Connexin 43 (Cx43), an astrocyte protein (also called Gap junction alpha-1 protein, GJA1), operates as an open pore via which toxic substances from the astrocytes reach the motor neurons. Patients with ALS who had a family history of the disease and those who had sporadic ALS had the most functional pores [5].

Connexin 31 (Cx31) shares amino acid and structural similarity to Cx43 [6, 7]. In their hexameric form, both connexins are composed of 6 subunits that work as transmembrane channels, binding together to form gap junction intercellular channels that open and close.

Hyperinsulinemia is a hallmark of T2D [8]. In the current study we performed molecular docking of insulin with the Cx43 monomer and Cx31 monomer. We then performed molecular docking of insulin with the Cx31 hexamer (also called Gap junction beta-3 protein, GJB3) to assess whether insulin might affect the pore.

## Methods

We used the program AutoDock Vina Extended for the docking study [9]. AutoDock Vina Extended achieves an approximately two orders of magnitude acceleration compared with the molecular docking software AutoDock 4, while also significantly improving the accuracy of the binding mode predictions. Further speed is achieved from parallelism, by using multithreading on multicore machines. AutoDock Vina Extended automatically calculates the grid maps and clusters the results in a way transparent to the user. UCSF Chimera 1.14 was used for molecular visualization [10].

The single chain human insulin molecule (humulin B) is from PubChem CID101896409. Human hexameric Cx31 hemichannel in the absence of calcium was deposited in the RCSB Protein Data Bank (6L3T) 2019-10-15, released: 2020-09-09.

Human insulin *in vivo* is a heterodimer of an A-chain and a B-chain, which are linked together by disulfide bonds. Heterodimeric human insulin was deposited in the RCSB Protein Data Bank (4EYN) 2012-5-01, released: 2013-05-01.

Cx43 had no entry in the RCBS Protein Data Bank for the complete molecule. We obtained a protein structure prediction for Cx43 (GJA1) from AlphaFold, an artificial intelligence (AI) system developed by Google’s DeepMind that predicts a protein’s 3D structure from its amino acid sequence. AlphaFold regularly achieves accuracy competitive with experiment [11, 12].

Cx31 had no entry in the RCBS Protein Data Bank for the monomer. We obtained a monomeric protein structure prediction for Cx31 (GJB3) from AlphaFold.

We used the ClusPro Server for protein-protein docking of human insulin (4EYN) to the Cx31 hexamer. ClusPro (https://cluspro.org) is a widely used tool for protein–protein docking. The server provides a simple home page for basic use, requiring only two files in Protein Data Bank (PDB) format [13]. The quality of automated docking by ClusPro is very close to that of the best human predictor groups [14].

We used GROMACS to perform molecular dynamics simulation of the human insulin (4EYN) docked to the Cx31 hexamer. GROMACS is a molecular dynamics package mainly designed for simulations of proteins, lipids, and nucleic acids.

## Results

Figure 1A shows Cx43 in AlphaFold, which produces a per-residue confidence score (pLDDT) between 0 and 100. Figure 1B displays predicted aligned error in Alpha Fold. Figure 1C shows CX31 in AlphaFold. Figure 1D displays Cx31 predicted aligned error in Alpha Fold.

**Figure 1A.**
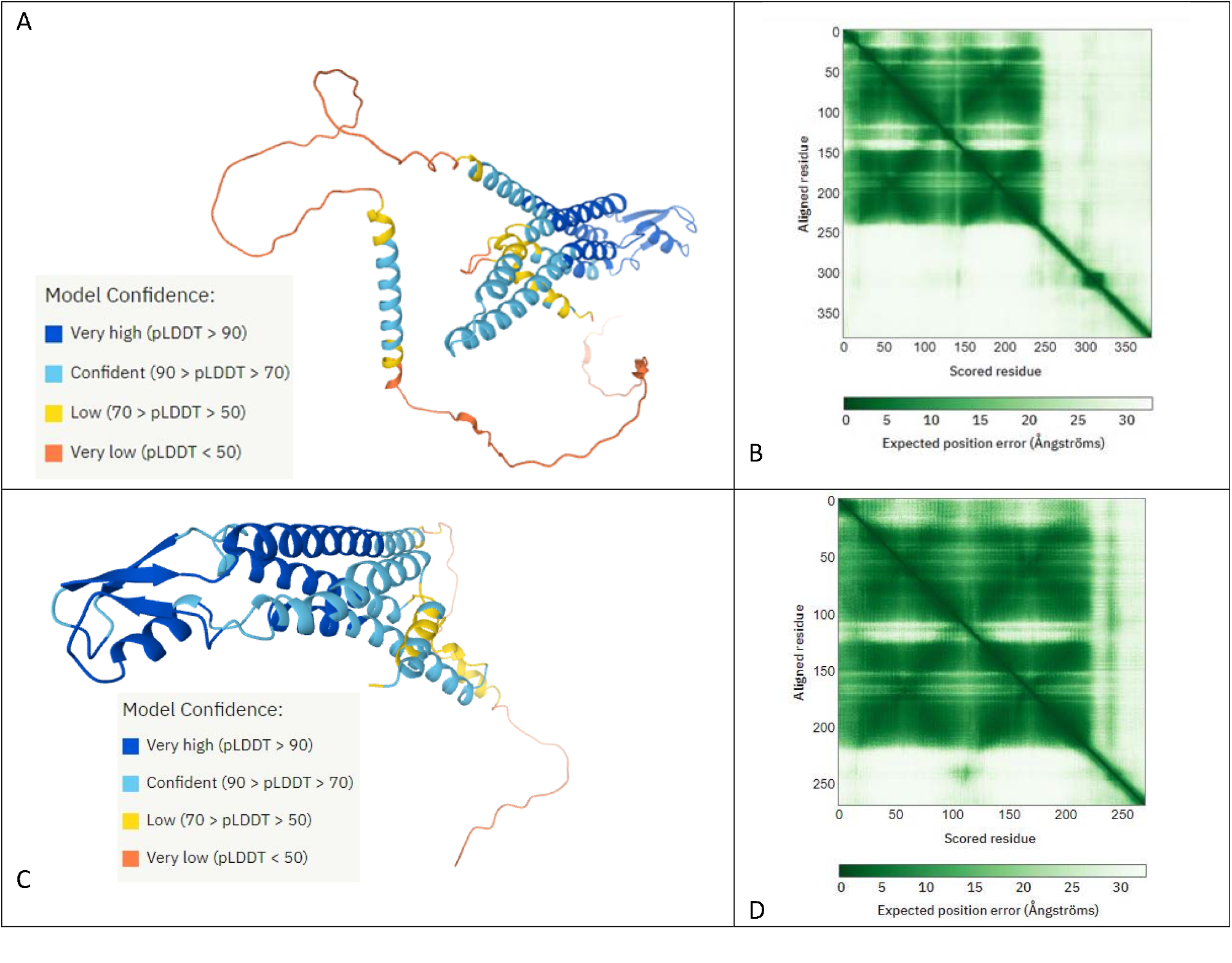
Cx43 (also called Gap junction alpha-1 protein, GJA1) in AlphaFold, which produces a per-residue confidence score (pLDDT) between 0 and 100 (lower left). Figure 1B. Predicted aligned error. The color at position (x, y) indicates AlphaFold’s expected position error at residue x, when the predicted and true structures are aligned on residue y. The shade of green indicates expected distance error in Ångströms. The color at (x, y) corresponds to the expected distance error in residue x’s position, when the prediction and true structure are aligned on residue y. Dark green is good (low error), light green is bad (high error). 1C. Cx31 (also called Gap junction beta-3 protein, GJB3) in AlphaFold. Figure 1D. Predicted Cx31 aligned error.

Figure 2A shows human Cx43 in its monomeric form docked with insulin. Arrows indicate the insulin molecule docked to the Cx43 N-terminal domain. Figure 2B is a closeup of the docked insulin molecule. Figure 2C shows human Cx31 in its monomeric form docked with insulin. Arrows indicate the insulin molecule docked to the Cx31 N-terminal domain. Figure 2D is a closeup of the docked insulin molecule. Note that insulin docks to almost the identical position in monomeric Cx31 and monomeric Cx43.

**Figure 2A.**
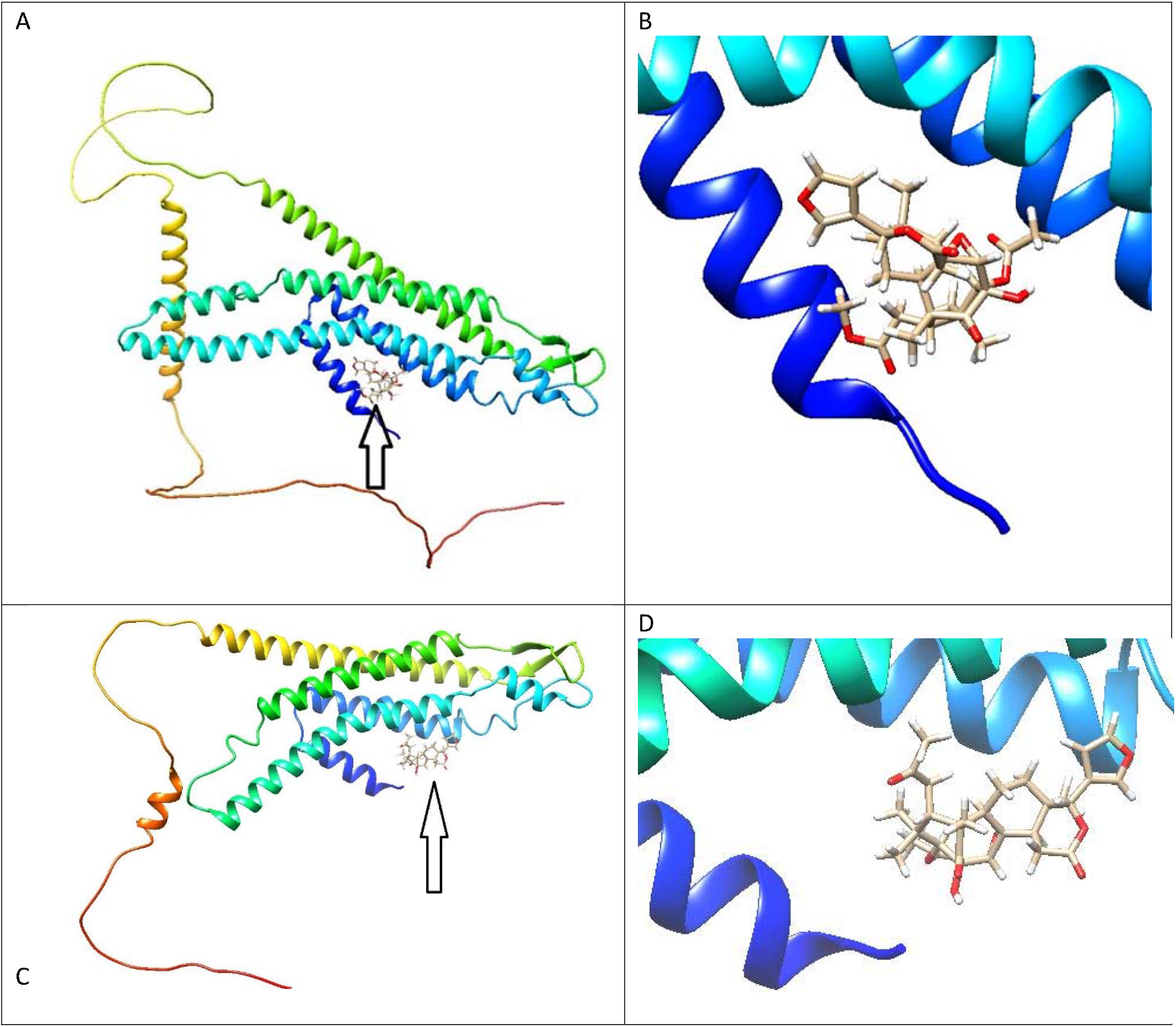
Human Cx43 in its monomeric form docked with humulin B single chain insulin molecule at the Cx43 N-terminal domain. Arrow indicates the insulin molecule. 2B. Closeup of Cx43 docked to insulin molecule. Per residue confidence score (pLDDT) from AlphaFold of the N-terminal domain (Figure 1A) is *confident*, 90 < pLDDT <70. 2C. Human Cx31 in its monomeric form docked with insulin at the Cx31 N-terminal domain. Arrow indicates the insulin molecule. 2D. Closeup of Cx31 docked to insulin molecule. Per residue confidence score (pLDDT) from AlphaFold of the N-terminal domain (Figure 1C) is *confident*, 90 < pLDDT <70. Note that insulin docks to almost the identical position in monomeric Cx31 and monomeric Cx43.

Figure 3 illustrates binding affinity (kcal/mol) calculated for 10 docking sites, insulin docked to monomeric Cx43. Only the one site with the highest affinity (upper left in figure) was a valid position.

**Figure 3.**
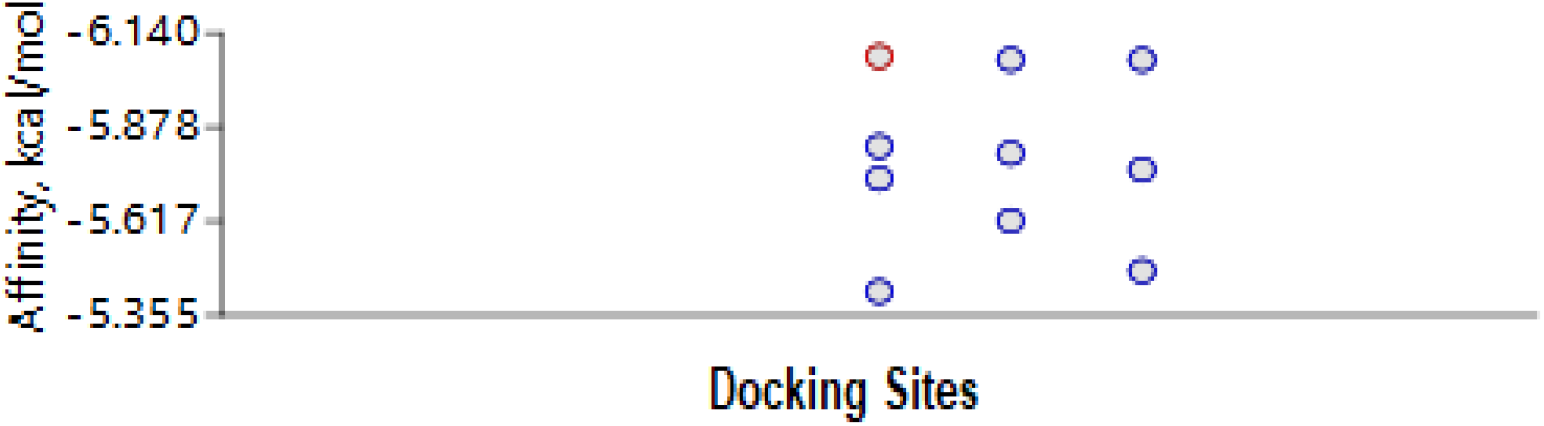
Binding affinity (kcal/mol) calculated for 10 docking sites, insulin to monomeric Cx43. Only one site with the highest affinity (upper left) was a valid position.

Table 1 contains docking parameters calculated by AutoDock Vina Extended, monomeric Cx43 to insulin. Lower values of root-mean-square deviations of atomic positions (RMSD) indicate that docking is validated with higher accuracy. RMSD values of 3 or more indicate no docking has occurred. One docking position, mode 1, with RMSD = 0 is highly valid.

**Table 1.** Docking parameters calculated by AutoDock Vina Extended for human monomeric Cx43 to insulin. Lower values of root-mean-square deviations of atomic positions (RMSD) indicate that docking is validated with higher accuracy. RMSD values of 3 or more indicate no docking has occurred. One docking position, mode 1, with RMSD = 0 is highly valid.

| mode | affinity (kcal/mol) | Ki(μmol) | RMSD lower bound(Å) | RMSD upper bound(Å) |
| --- | --- | --- | --- | --- |
| 1 | -6.074602412 | 35.26029 | 0 | 0 |
| 2 | -6.065978959 | 35.77725 | 27.73112185 | 31.15029382 |
| 3 | -6.065369451 | 35.81407 | 2.902005898 | 8.116629233 |
| 4 | -5.824558059 | 53.77367 | 1.92658046 | 7.437705569 |
| 5 | -5.805591846 | 55.52288 | 2.733718603 | 8.206586658 |
| 6 | -5.759997329 | 59.96433 | 2.771359767 | 8.194987559 |
| 7 | -5.735407323 | 62.50541 | 2.208205522 | 5.237036912 |
| 8 | -5.617848751 | 76.22346 | 2.149206141 | 7.494382604 |
| 9 | -5.477866589 | 96.53755 | 15.877928 | 19.02405307 |
| 10 | -5.420853417 | 106.2887 | 2.428831289 | 5.403085509 |

Figure 4 shows binding affinity (kcal/mol) calculated for 10 docking sites, insulin to monomeric Cx31. Only one site with the highest affinity (upper left) was a valid position.

**Figure 4.**
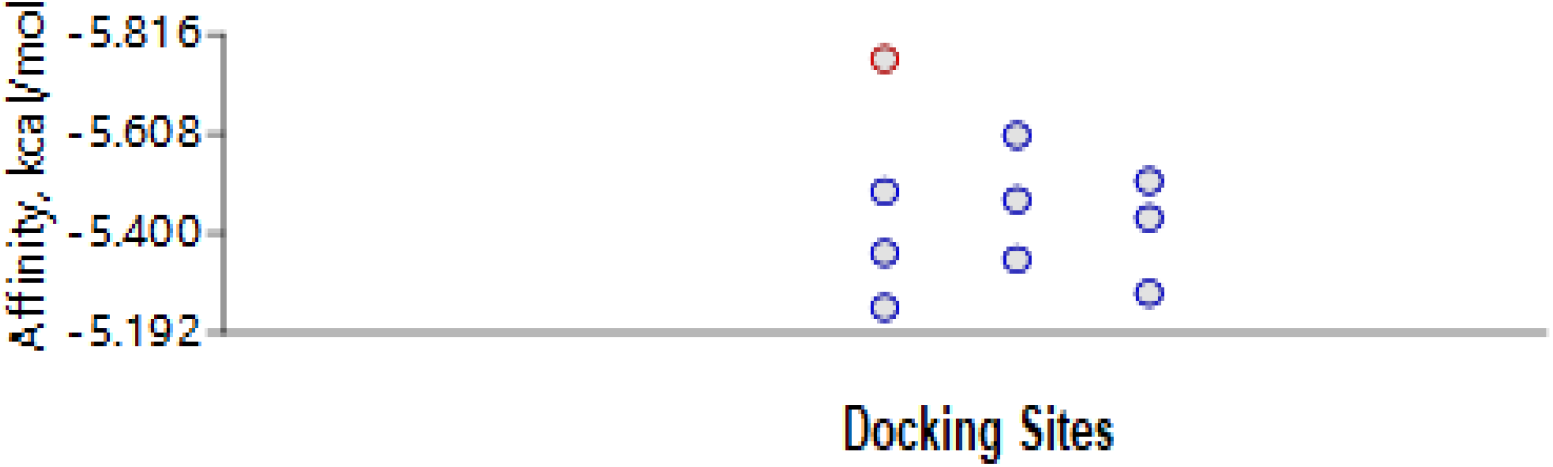
Binding affinity (kcal/mol) calculated for 10 docking sites, insulin to monomeric Cx31. Only one site with the highest affinity (upper left) was a valid position.

Table 2 contains docking parameters calculated by AutoDock Vina Extended, monomeric Cx31 to insulin. One docking position, mode 1, with RMSD = 0 is highly valid.

**Table 2.** Docking parameters calculated by AutoDock Vina Extended for human monomeric Cx31 to insulin. One docking position, mode 1, with RMSD = 0 is highly valid.

| mode | affinity (kcal/mol) | Ki(μmol) | RMSD lower bound($\text{\AA}$) | RMSD upper bound($\text{\AA}$) |
| --- | --- | --- | --- | --- |
| 1 | -5.764226326 | 59.53785 | 0 | 0 |
| 2 | -5.604354028 | 77.97948 | 20.29215496 | 24.5788459 |
| 3 | -5.50895568 | 91.6026 | 24.13479769 | 26.64546373 |
| 4 | -5.487040562 | 95.05428 | 24.10361311 | 28.26772351 |
| 5 | -5.469198364 | 97.9603 | 25.81208942 | 28.4784468 |
| 6 | -5.431554356 | 104.3863 | 19.49592317 | 23.8544476 |
| 7 | -5.358509811 | 118.0825 | 3.708488437 | 4.594993627 |
| 8 | -5.344926538 | 120.8209 | 24.15928651 | 26.64988346 |
| 9 | -5.274514921 | 136.0674 | 25.04492059 | 29.83058177 |
| 10 | -5.244106707 | 143.2331 | 25.62035767 | 28.69788407 |

Figure 5 illustrates binding affinity (kcal/mol) calculated for 10 docking sites, insulin docked to hexameric Cx31. Only the one site in the hemichannel with the highest affinity (upper left in figure) was a valid position.

**Figure 5.**
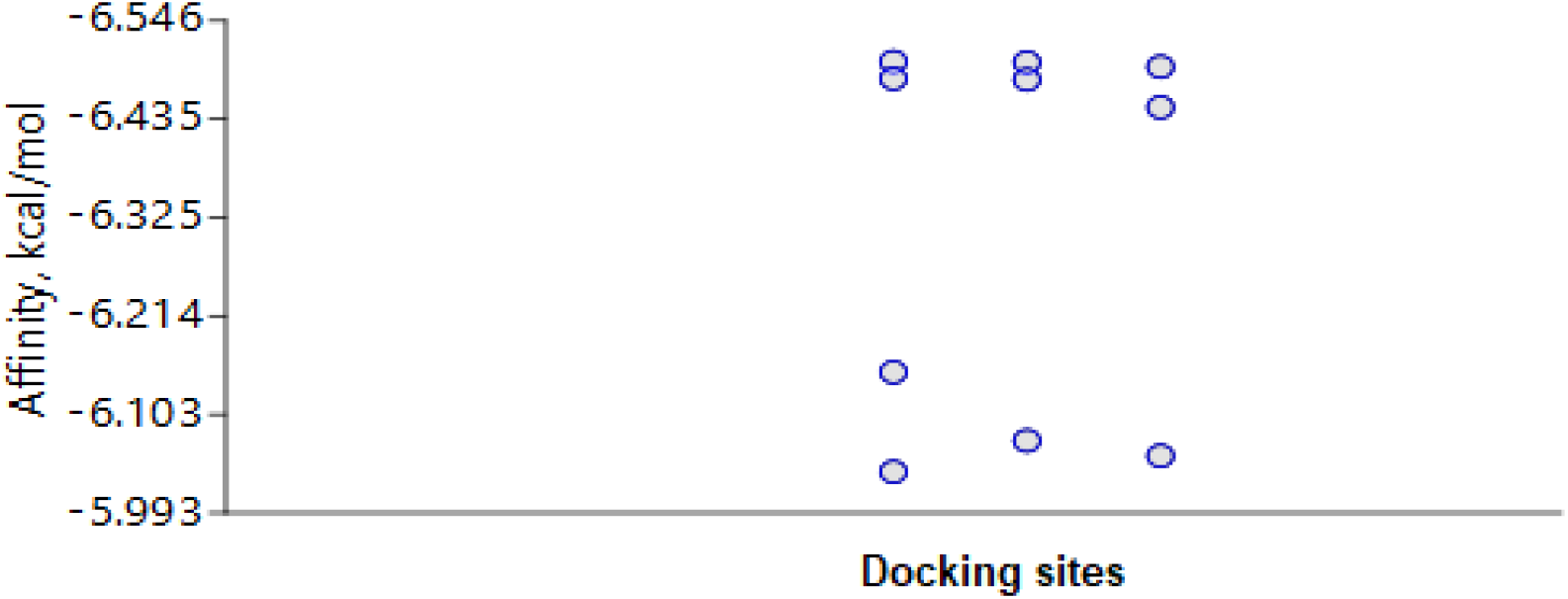
Binding affinity (kcal/mol) calculated for 10 docking sites, humulin B insulin to Cx31 hexamer. Only one site with the highest affinity (upper left) was a valid position.

Figure 6A shows human Cx31 in its hexameric form docked with insulin. Arrows indicate the insulin molecule docked within the hemichannel. Figure BB is a closeup of the docked insulin molecule.

**Figure 6A.**
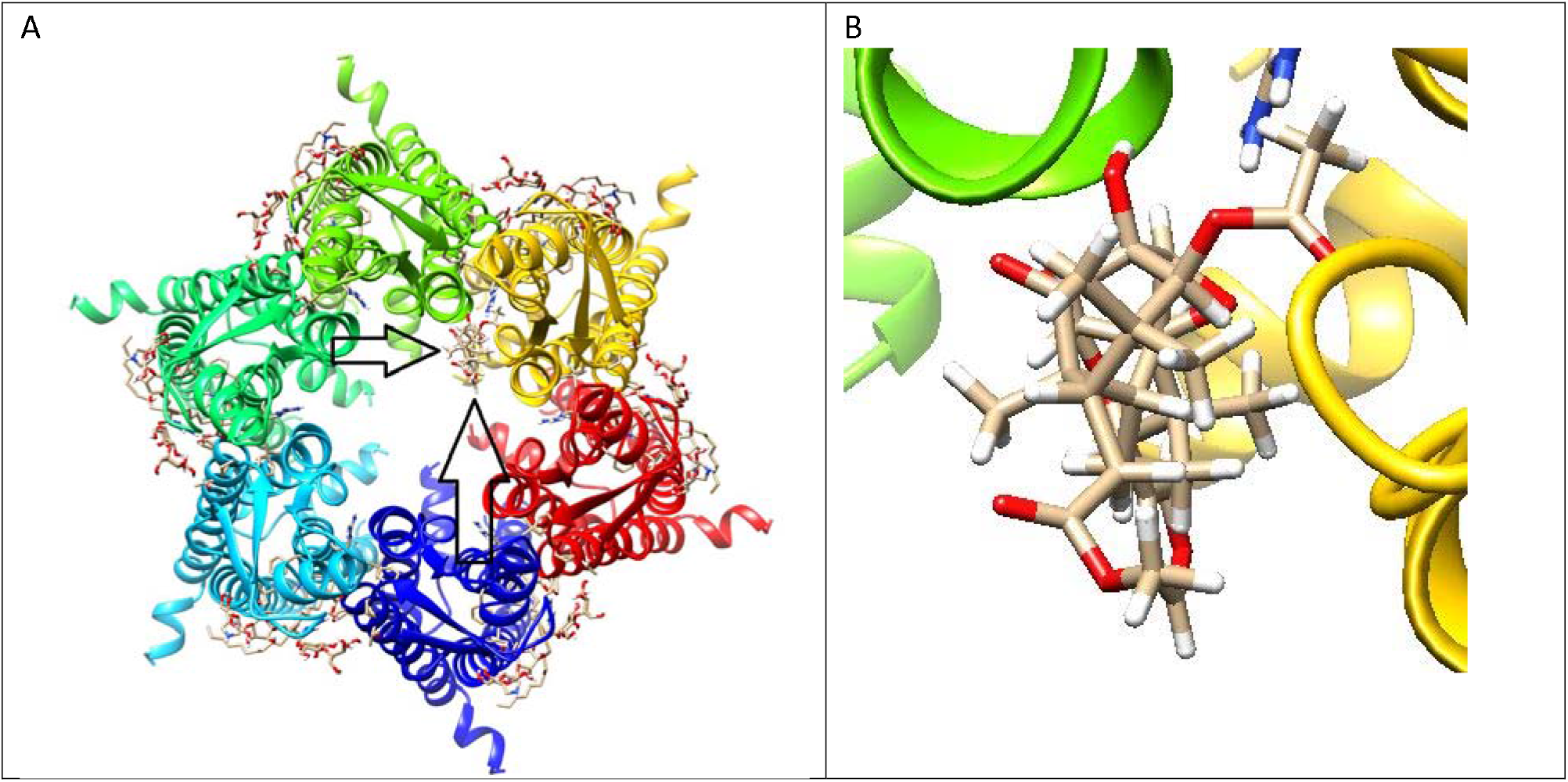
Human Cx31 in its hexameric form with open hemichannel (center) docked with humulin B single chain insulin molecule. Arrows indicate the insulin molecule within the hemichannel. 6B. Closeup of docked humulin B, single chain insulin molecule.

Table 3 contains docking parameters calculated by AutoDock Vina Extended, hexameric Cx31 to insulin. One docking position, mode 1, with RMSD = 0 is highly valid.

**Table 3.** Docking parameters calculated by AutoDock Vina Extended of hexameric Cx31 to insulin. One docking position, mode 1, with RMSD = 0 is highly valid.

| mode | affinity (kcal/mol) | Ki(μmol) | RMSD lower bound($\text{\AA}$) | RMSD upper bound($\text{\AA}$) |
| --- | --- | --- | --- | --- |
| 1 | -6.499765043 | 17.2044 | 0 | 0 |
| 2 | -6.498996353 | 17.22674 | 8.738951896 | 10.75563818 |
| 3 | -6.49403788 | 17.37151 | 16.04060202 | 18.62194177 |
| 4 | -6.480787951 | 17.76437 | 8.710319846 | 10.73125405 |
| 5 | -6.479632275 | 17.79906 | 18.41293941 | 21.47099771 |
| 6 | -6.448499272 | 18.75934 | 16.05073372 | 18.60996308 |
| 7 | -6.150886428 | 31.00055 | 28.43659168 | 33.35036486 |
| 8 | -6.073789675 | 35.30869 | 22.76797584 | 26.71361825 |
| 9 | -6.056759246 | 36.33834 | 34.99424228 | 38.79479813 |
| 10 | -6.038860805 | 37.45283 | 40.62664359 | 44.9530327 |

Figure 7A. Human insulin (4EYN), the heterodimer, A-chain (left) and B-chain (right). 7B. Human Cx31 in its hexameric form with open hemichannel (center) docked with human insulin by the ClusPro Server. Note that the human insulin heterodimer completely blocks the open hemichannel.

**Figure 7A.**
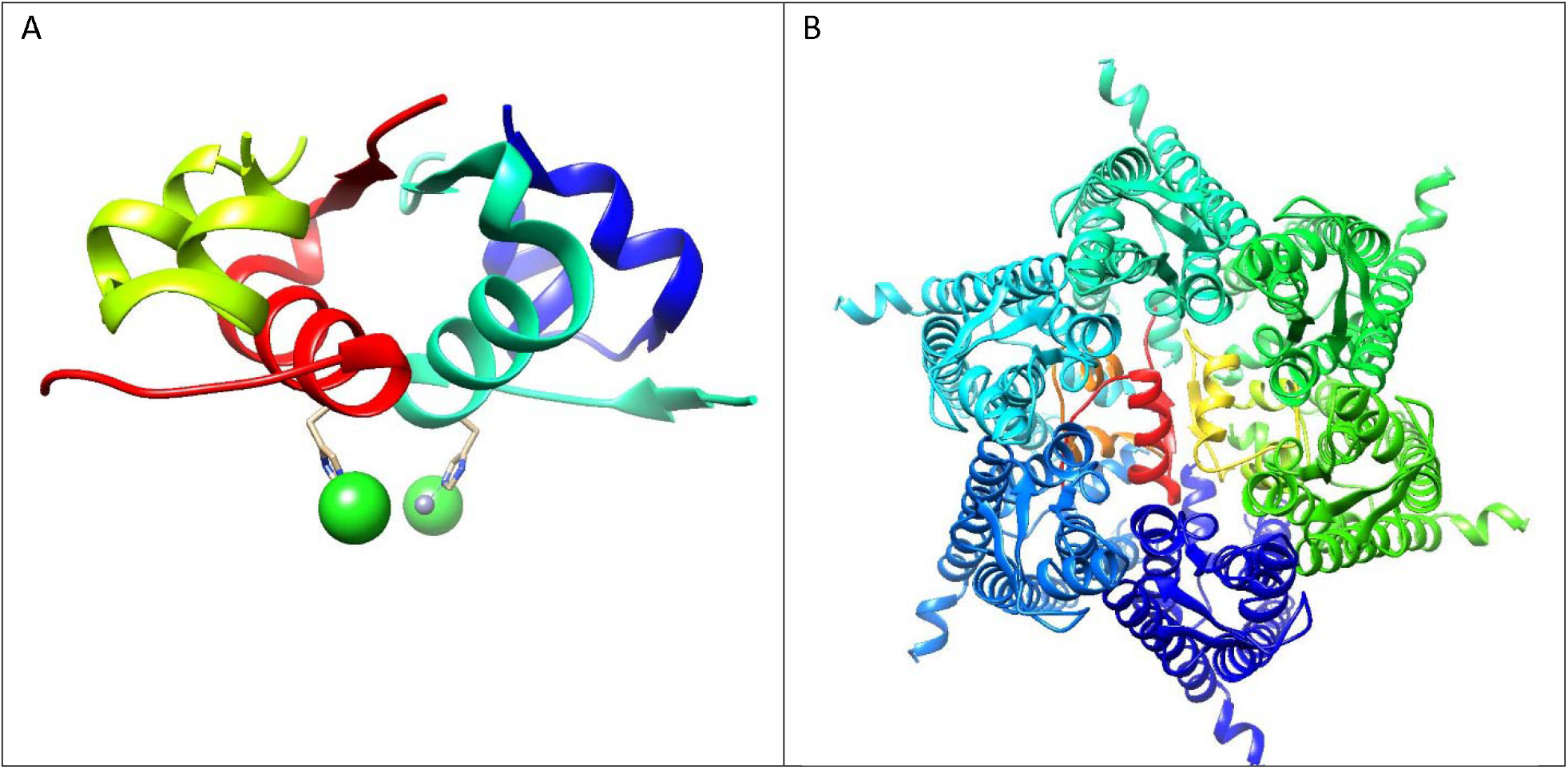
Human insulin (RCBS Protein Data Bank 4EYN), the heterodimer, A-chain (left) and B-chain (right). 7B. Human Cx31 in its hexameric form with open hemichannel (center) docked with human insulin in highest ranked ClusPro configuration. Note that the human insulin heterodimer completely blocks the open hemichannel (center).

Figure 8 shows the first six configurations (clusters) of Human Cx31 in its hexameric form, open hemichannel docked with human insulin, results from ClusPro. Configuration 0, the highest ranked, is shown enlarged and rotated in figure 7B. Note that the human insulin heterodimer (red) is in the identical position in all six configurations, blocking the open Cx31 hemichannel.

**Figure 8.**
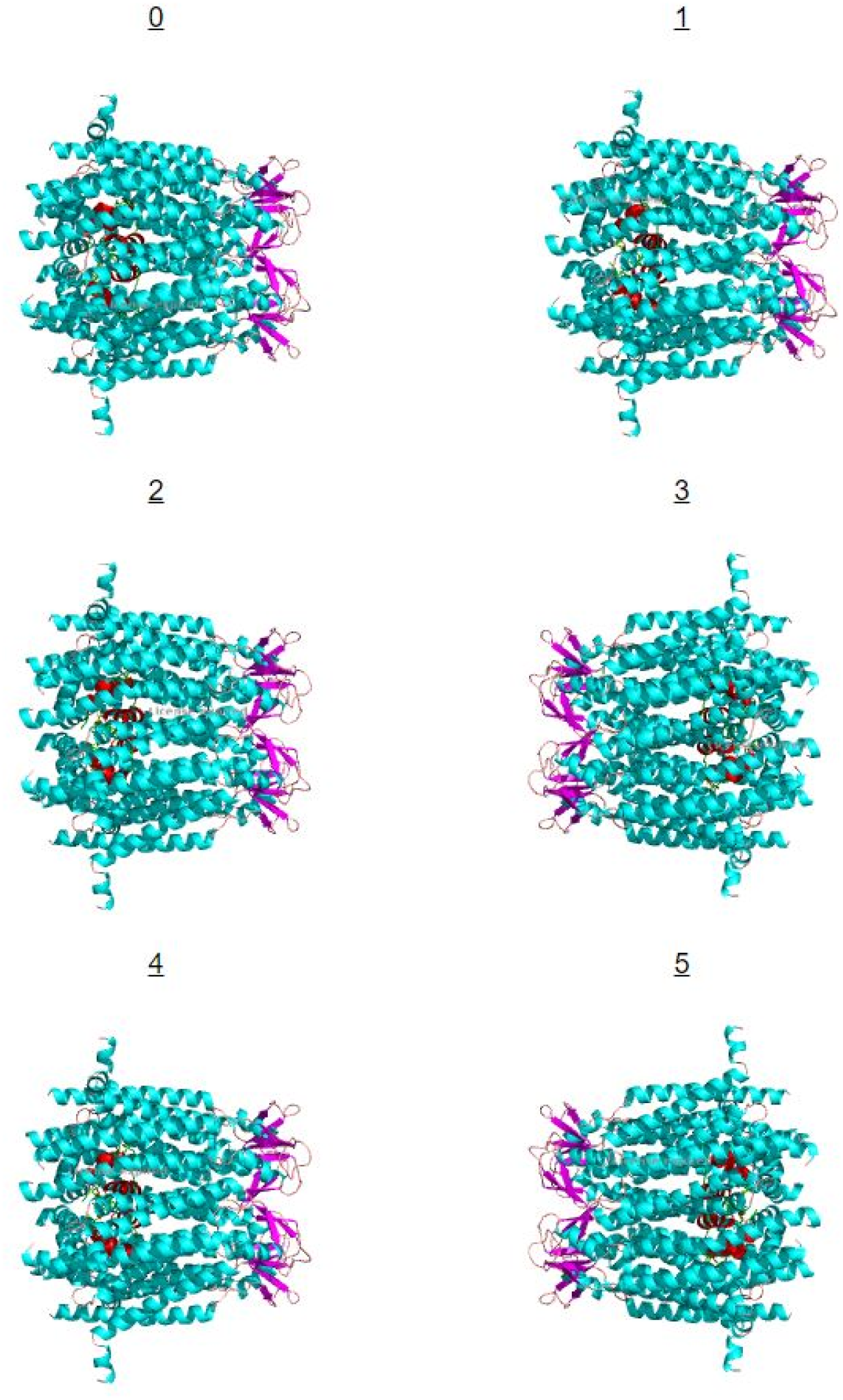
First six configurations (clusters) of Human Cx31 in its hexameric form, open hemichannel docked with human insulin, results from ClusPro. Configuration 0, the highest ranked, is shown enlarged and rotated in figure 7B. Note that the human insulin heterodimer (red) is in the identical position in all six configurations, blocking the open Cx31 hemichannel.

Table 4 lists cluster scores and energies for Human Cx31 in its hexameric form with open hemichannel docked with human insulin heterodimer. The ligand position with the most *neighbors* within 9 angstroms becomes a cluster center, and its neighbors the members of the cluster. These are then removed from the set and a second cluster center is located, then a third, up to cluster 5. Thus, the cluster rank is determined.

**Table 4.**
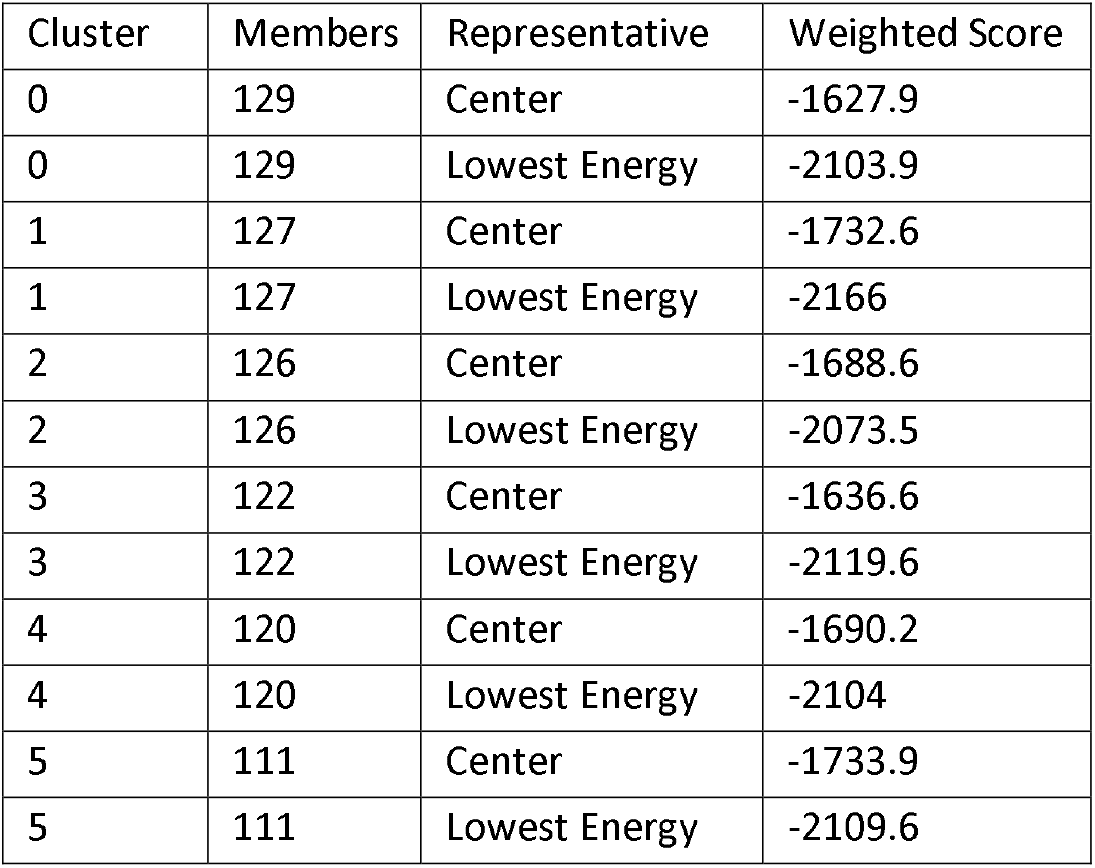
Cluster scores and energies for Human Cx31 in its hexameric form with open hemichannel docked with human insulin heterodimer. The ligand position with the most *neighbors* within 9 angstroms becomes a cluster center, and its neighbors the members of the cluster. These are then removed from the set and a second cluster center is located, then a third, up to cluster 5. Thus, the cluster rank is determined.

| Cluster | Members | Representative | Weighted Score |
| --- | --- | --- | --- |
| 0 | 129 | Center | -1627.9 |
| 0 | 129 | Lowest Energy | -2103.9 |
| 1 | 127 | Center | -1732.6 |
| 1 | 127 | Lowest Energy | -2166 |
| 2 | 126 | Center | -1688.6 |
| 2 | 126 | Lowest Energy | -2073.5 |
| 3 | 122 | Center | -1636.6 |
| 3 | 122 | Lowest Energy | -2119.6 |
| 4 | 120 | Center | -1690.2 |
| 4 | 120 | Lowest Energy | -2104 |
| 5 | 111 | Center | -1733.9 |
| 5 | 111 | Lowest Energy | -2109.6 |

Figure 9 is a molecular dynamics simulation of human Cx31 in its hexameric form with open hemichannel docked with human insulin heterodimer. Time series shows the RMSD levels fluctuation ∼ 0.1 nm (1 Å), indicating that the structure is quite stable. The reasonably invariant radius of gyration (Rg) values indicate that the docked protein remains highly stable over the course of 1 ns. These results suggest that once insulin blocks the Cx31 open hemichannel, the block is stable and will remain in place.

**Figure 9 A.**
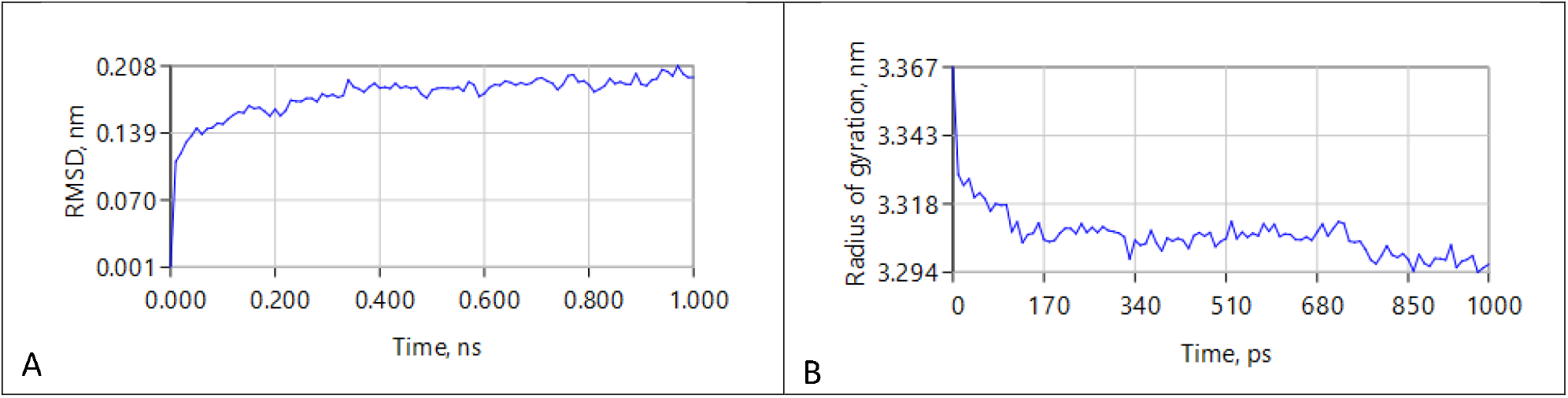
Molecular dynamics simulation human Cx31 in its hexameric form with open hemichannel docked with human insulin heterodimer. Time series shows the RMSD levels fluctuation of ∼ 0.1 nm (1 Å), indicating that the structure is quite stable. 9B. Radius of gyration (Rg). The reasonably invariant Rg values indicate that the docked protein remains highly stable over the course of 1 ns. These results suggest that once insulin blocks the Cx31 open hemichannel, the block is stable and will remain in place.

## Discussion

Cx43 is normally expressed in astrocytes and found in most human astrocytomas and in the astroglial component of glioneuronal tumors [15]. Astrocytes may cause the death of motor neurons since astrocyte dysfunction occurs after symptom onset in patients with ALS. Astrocytes provoke motor neuron death through Cx43 hemichannels that allow the movement of toxic astrocyte substances into motor neurons. Cx43 protein is elevated in cerebrospinal fluid of ALS patients and could be a biomarker [5].

Because astrocytes may be responsible for the progression of ALS, drugs that block Cx43 hemichannels might be therapeutic [5]. One analysis of the Cx43 structure revealed a closed sieve-like molecular gate [16]. The Cx43 N-terminal domain is involved in channel gating and oligomerization and may control the switch between the channel’s open and closed states. Therefore, the docking of insulin to the Cx43 N-terminal domain we demonstrate here may be capable of blocking the channel, preventing toxic astrocyte substances from entering and destroying motor neurons.

Our finding that insulin docks within the open hemichannel of hexameric Cx31, potentially blocking it, again suggests that the block may be responsible for the inverse relationship between ALS and T2D. This conclusion is reinforced by the fact that insulin docks to the same position, the N-terminal domain, of monomeric Cx31 and monomeric Cx43 (Figure 2). Molecular dynamics simulation indicates that once insulin blocks the Cx31 open hemichannel, the block is stable and will remain in place. Therefore, insulin might be a treatment for ALS, especially since insulin enhances glucose-stimulated insulin secretion in healthy humans [17]. Type 1 diabetes, characterized by a total lack of insulin, is associated with increased risk of ALS [1, 2], again suggesting that insulin could have a beneficial effect on ALS.

Therapeutic insulin would likely need to be administered intranasally, rather than parenterally, for an adequate dose to the cerebrospinal fluid (CSF). CSF insulin levels are normally much lower than plasma insulin levels [18]. Normal plasma insulin is 50 pmol/l compared to normal CSF insulin, 3.3 pmol/l [19]. Intranasal insulin achieves direct access to the CSF within 30 minutes, bypassing the bloodstream. Insulin secretogogues such as oral sulfonylurea or glinide might also be of value.

However, this hypothesis has weaknesses:

1. Zhang et al did not observe a significant association between 2 h glucose, fasting glucose, fasting insulin, fasting proinsulin, HbA1C and the risk of ALS in European populations [4]. But Zhang et al measured fluctuations in plasma insulin that may have been related to much smaller changes in CSF insulin, given the low CSF insulin levels normally observed [19]. Moreover, the Cx43 gene, GJA1, has multiple polymorphisms, and some may affect Cx43 interaction with insulin [20].
2. Insulin docks almost identically within monomeric Cx31 and monomeric Cx43. We assume that insulin docks identically within hexameric Cx31 and hexameric Cx43, but this may not be the case. Cx31 abnormalities are associated with erythrokeratoderma, a rare skin condition [21].

In sum, the docking results presented here indicate that insulin may be capable of inhibiting the progression of ALS by blocking Cx43 hemichannels. Further study is warranted.

